# A cryptic local genetic cluster in Northern France amid the European mosaic of flat oyster lineages revealed by integrating SNP array and whole-genome sequencing

**DOI:** 10.64898/2026.06.26.734753

**Authors:** Sylvie Lapègue, Florence Cornette, Serge Heurtebise, Stéphane Pouvreau, Cynthia Carpentier, Lila Colston-Népali, Nicolas Bierne, Céline Reisser

**Affiliations:** MARBEC, Univ Montpellier, CNRS, Ifremer, IRD, Montpellier, France; MARBEC, Univ Montpellier, CNRS, Ifremer, IRD, Sète, France; Ifremer, ASIM, La Tremblade, France; Ifremer, LEMAR, Plouzané, France; CAPENA, Le Château d’Oléron, France; ISEM, Univ Montpellier, CNRS, IRD, Montpellier, France

**Keywords:** *Ostrea edulis*, SNP array, Whole genome sequencing, cryptic lineage, restoration

## Abstract

The European flat oyster (*Ostrea edulis*), like numerous other oyster species, has been exploited for millennia and cultivated and translocated for centuries. Following a severe population decline, and in the context of ongoing conservation and restoration programs, genetic considerations must now be addressed to avoid mistakes. The objective of our study was to complement population genetic studies conducted at various scales along European coasts. Our sampling primarily targeted the French Atlantic, English Channel, and Mediterranean coasts, aiming to provide a fine-scale genetic characterization of populations in these regions. By integrating SNP array and low-coverage sequencing datasets, we obtained a comprehensive overview of the population genetic structure of *Ostrea edulis* across western Europe. Most previously identified clusters in Western Europe were confirmed. In France, populations assigned to these clusters exhibited notable within-patch homogeneity. However, two key findings emerged: (1) an extensive overlap zone between the Atlantic and western Mediterranean clusters, spanning at least from southern Portugal to southern France, and (2) the detection of a novel, clearly distinct cryptic cluster east of the English Channel, whose geographic range remains to be better delineated. These insights are critical for informing management decisions, particularly as restoration and conservation plans are currently being implemented across the species’ range.

## Introduction

Oysters have been exploited for millennia and cultivated and translocated for centuries (Neild 1995). Today, they are the most widely cultivated molluscan species (Botta et al. 2020). Beyond their economic importance, oyster reefs serve as vital estuarine ecosystems, supporting rich and diverse populations of invertebrates and fish (Cole et al. 2022). However, oyster reefs have suffered the greatest loss of all marine ecosystems, with an estimated 85% global decline (Beck et al. 2011). In some bays and estuaries worldwide, oyster populations have declined by 90–99%, even reaching a state of functional extinction (McAfee and Connell 2021; Smith et al. 2023).

Until recently, the past distribution and extent of oyster reefs remained unknown. Several studies focusing on historical record analyses have since provided valuable insights for the current management of these habitats (zu Ermgassen et al. 2012). Ancient or degraded reefs may thus serve as starting points for restoration programs (Smith et al. 2023). While early oyster restoration efforts primarily aimed to rebuild oyster production for harvest, many contemporary projects have shifted their focus toward recovering ecosystem services (Coen and Luckenbach 2000). These services include water filtration, benthic–pelagic coupling, nutrient cycling, sediment stabilization, and habitat provision.

The Eastern oyster, *Crassostrea virginica*, was the first oyster species targeted by restoration projects, which expanded significantly in the 1990s and 2000s (Duarte et al. 2020). A synthesis of 1,768 projects conducted in the United States since 1964 revealed that oyster substrate restoration efforts have been concentrated in the Chesapeake Bay and the Gulf Coast, re-establishing 4.5% of the lost reef area in these regions (Bersoza Hernandez et al. 2018). The Billion Oyster Project, launched in the 2010s, aims to restore one billion oysters in New York Harbor by 2035, making it one of the most ambitious restoration initiatives worldwide (https://www.billionoysterproject.org). Restoration efforts for other oyster species, including *Ostrea lurida*, *Ostrea edulis*, *Ostrea angasi*, and *Saccostrea glomerata*, increased in the 2010s and continue to expand in the 2020s (Smith and Pruett 2025).

The European flat oyster, *Ostrea edulis*, was once a common species across Europe (Thurstan et al. 2024). Due to overharvesting in the late 19th century and disease outbreaks in the 20th century (Pouvreau et al. 2023), it is now classified as “Threatened” or “Declining” by the Convention for the Protection of the Marine Environment of the North-East Atlantic (OSPAR 2008). Under the IUCN (International Union for Conservation of Nature) criteria, its reefs could even be considered “Collapsed” (zu Ermgassen et al. 2025). Nevertheless, significant restoration efforts have been undertaken in Europe over the past two decades. In 2026, nearly 50 projects focused on European flat oyster bed restoration are listed by the NORA Europe network (noraeurope.eu).

To avoid failure, restoration handbooks are available (e.g., zu Ermgassen et al. 2021), but genetic considerations are rarely addressed. Although the Berlin Oyster Recommendations emphasize the importance of “preserving the extant genetic diversity of native oysters in Europe” (Pogoda et al. 2019), the genetic characteristics of wild populations and introduced stocks are seldom considered at the outset of restoration programs. Depending on the level of reef depletion, several options are available for active oyster reef restoration (Pouvreau et al. 2023). For instance, when larval abundance in the wild remains sufficient, facilitating successful settlement is an effective and efficient option, particularly when implemented with minimal environmental impact (Kamermans et al. 2025). However, when populations are severely depleted, the introduction of adult oysters as broodstock may be considered (Colsoul et al. 2021; Sas et al. 2020). Regardless of the approach, characterizing local populations and introduced stocks is critical. This includes assessing their genetic characteristics, such as local genetic diversity and genetic relationships among populations or stocks. Understanding how adaptive and quantitative genetic variation is distributed within and among populations is essential, as restoration efforts could inadvertently introduce outbreeding depression when locally adapted and potentially non-adapted animals are brought into contact (Camara and Vadopalas 2009; Laikre et al. 2010).

In Europe, genetic studies of the European flat oyster, *O. edulis*, began 40 years ago with allozyme data (Blanc et al. 1986). Early studies using these markers (Saavedra et al. 1995), followed by analyses of mitochondrial DNA (Díaz-Almela et al. 2004) and microsatellite markers (Launey et al. 2002), revealed weak but significant genetic differentiation between Atlantic and Mediterranean populations, apparently consistent with an isolation-by-distance model. Subsequent studies using additional microsatellite markers (Vera et al. 2016) and hundreds (Lapègue et al. 2023) or thousands (Vera et al. 2019) of SNPs identified four geographically distinct genetic clusters across Europe. However, this global pattern may have been locally obscured by historical translocations, which were frequently performed between European countries during the last century (Bromley et al. 2016). For example, such translocations were suspected from the Atlantic coast to the Venice Lagoon (Lapègue et al. 2023), and genomic evidence of past translocations has been observed from Norway to Sweden and Denmark (Alves Monteiro et al. 2024). The shift toward genomic approaches, combined with fine-scale sampling, has enabled the detection of a finer population structure, including additional clusters in Scandinavia. Nevertheless, these findings remain coherent with the broader European genetic pattern (Alves Monteiro et al. 2024; Robert et al. 2025).

Three putative chromosomal inversions were identified in *O. edulis* (Lapègue et al. 2023; Alves Monteiro et al. 2024, Valk et al. 2026a) with a potential fourth (Ewers et al., 2025). These inversions, ancient and likely under divergent selection, show high allele frequency differences between a northern lineage in Scandinavian and North Sea populations and Southern lineage in Atlantic and Mediterranean ones (Alves Monteiro et al., 2024). Additionally, two of these inversions exhibit parallel differentiation signals, a pattern also observed in some non-inverted genomic regions (“islands of differentiation”), suggesting long-term barriers to gene flow (Colston-Nepali, 2025). These findings, combined with secondary contact signatures, support the existence of cryptic, semi-isolated lineages within the species, with a shared evolutionary history partially erased by gene flow (Lapègue et al., 2023; Colston-Nepali, 2025).

In this study, we focused on the geographical patterns retained by the majority of the genome, excluding known inversions. Our sampling primarily targeted the French Atlantic, English Channel, and Mediterranean coasts, with the aim of providing a fine-scale genetic characterization of populations in these regions, where numerous restoration and conservation programs are currently being implemented (e.g., FOREVER: [Pouvreau et al. 2021]; REFONA [Carpentier et al., 2024]). Previous large-scale studies included only a limited number of samples from France, typically from Brittany (Atlantic coast) and Occitanie (Mediterranean coast), which precluded any exploration of regional genetic structure (Lapègue et al., 2023; Alves Monteiro et al., 2024). To place our regional findings within a broader European context, we integrated genotypes from a medium-density marker array (Gutierrez et al., 2017) with whole-genome sequencing (WGS) data from European reference populations (Alves Monteiro et al., 2024). Our objectives were twofold: first, to expand French sampling to clarify genetic patterns at the national scale; and second, to combine our SNP array data with low-coverage WGS datasets, particularly to account for northern European clusters that were not represented in our sampling. This approach enabled us to identify a previously undetected cryptic genetic cluster and to better characterize the contact zones between clusters present in France. Collectively, these findings provide critical insights for managers involved in the design and implementation of conservation and restoration projects for this species/habitat along the French marine coasts.

## Materials and Methods

### Biological material and DNA extraction

A total of 20 populations of *O. edulis* were sampled along the French coast of the English Channel, Atlantic Ocean and Mediterranean Sea from 2019 to 2023. The number of samples per population varied from 7 to 72 for a total number of 487 oysters (Table 1). Genomic DNA was extracted from gill, mantle or muscle tissue, using the QIAamp DNA mini-kit (QIAGEN) according to the manufacturer’s recommendations. These samples are referred to as the Genotyped Populations (GP).

**Table 1.** Location of sampling populations included in this study. Population identification codes (ID) in bold for Genotyped Populations (GP), and in italics for Sequenced Populations (SP). N = Number of samples. *Ho* = Observed Heterozygote estimates for Merged Dataset II

| Country | Location | ID | Latitude | Longitude | N | *Ho* | Missing data (%) | Sampling Year | Original study |
| --- | --- | --- | --- | --- | --- | --- | --- | --- | --- |
| Norway | Ostretjonn | *OSTR* | 58,32 | 6,25 | 19 | 0.212 | 13.5 | 2020 | Alves Monteiro et al. 2024 |
| Norway | Aga Bomlo | *AGAB* | 59,84 | 5,24 | 19 | 0.219 | 14 | 2020 | Alves Monteiro et al. 2024 |
| Norway | Haugevagen | *HAUG* | 59,29 | 5,19 | 21 | 0.214 | 15.8 | 2020 | Alves Monteiro et al. 2024 |
| Norway | Bunnefjorden | *BUNN* | 59,74 | 10,72 | 18 | 0.211 | 21.4 | 2021 | Alves Monteiro et al. 2024 |
| Sweeden | Hyppeln | *HYPP* | 57,76 | 11,61 | 19 | 0.212 | 22.4 | 2020 | Alves Monteiro et al. 2024 |
| Denmark | Logstor | *LOGS* | 56,97 | 9,14 | 19 | 0.225 | 12.5 | 2020 | Alves Monteiro et al. 2024 |
| Netherlands | Wadden Sea | *WADD* | 53,49 | 6,59 | 16 | 0.225 | 15.2 | 2020 | Alves Monteiro et al. 2024 |
| Netherlands | Lake Grevelingen | *GREV* | 51,76 | 3,91 | 16 | 0.205 | 16.4 | 2020 | Alves Monteiro et al. 2024 |
| Scotland | Lochryan | *RYAN* | 54,95 | -5,03 | 20 | 0.200 | 15.8 | 2020 | Alves Monteiro et al. 2024 |
| Ireland | Tralee Bay | *TRAL* | 52,29 | -9,93 | 18 | 0.195 | 14.6 | 2013 | Alves Monteiro et al. 2024 |
| England | Tollesbury | *TOLL* | 51,75 | 1,16 | 19 | 0.196 | 18.5 | 2020 | Alves Monteiro et al. 2024 |
| France | Bois de Cise | **CIS** | 50,09 | 1,42 | 11 | 0.243 | 0.1 | 2022 | This study |
| France | Manche Est | **MAN** | 50,09 | 1,27 | 29 | 0.239 | 0.1 | 2022 | This study |
| France | Granville | **GRAN** | 50,09 | 1,42 | 26 | 0.230 | 0.1 | 2012 | This study |
| France | La Rance | **R** | 48,55 | -1,97 | 20 | 0.231 | 0.9 | 2009 | This study |
| France | Pomelin | **POME** | 48,76 | -3,12 | 25 | 0.224 | 0.1 | 2020 | This study |
| France | Touil An Houillet | **TOUL** | 48,76 | -3,12 | 26 | 0.230 | 0.1 | 2020 | This study |
| France | Brest | **BZH** | 48,32 | -4,35 | 9 | 0.226 | 0.1 | 2020 | This study |
| France | Keraliou | **LIO** | 48,37 | -4,43 | 26 | 0.242 | 1.8 | 2020 | This study |
| France | Roz | **LEF** | 48,32 | -4,34 | 27 | 0.231 | 3.8 | 2020 | This study |
| France | Pont L'Abbée | **PLAB** | 47,85 | -4,18 | 26 | 0.225 | 0.1 | 2020 | This study |
| France | Odet | **ODET** | 47,92 | -4,46 | 25 | 0.231 | 0.1 | 2020 | This study |
| France | Ria d'Etel | **ETEL** | 47,69 | -3,16 | 26 | 0.232 | 0.1 | 2020 | This study |
| France | Quiberon | **BQU** | 47,57 | -3,08 | 26 | 0.229 | 0.2 | 2020 | This study |
| France | Baie de Bourgneuf | **BOUR** | 47,02 | -2,11 | 21 | 0.227 | 2.8 | 2020 | This study |
| France | Petite Chette | **CHET** | 45,94 | -1,20 | 72 | 0.229 | 0.2 | 2022-2023 | This study |
| France | Arcachon | **ARC** | 44,59 | -1,23 | 49 | 0.239 | 0.8 | 2022-2023 | This study |
| Spain | Pontedeume | *PONT* | 43,41 | -8,18 | 16 | 0.190 | 18.9 | 2013 | Alves Monteiro et al. 2024 |
| France | Palavas | **PAL** | 43,52 | 3,91 | 8 | 0.216 | 0.2 | 2020 | This study |
| France | Toulon | **TOU** | 43,11 | 5,90 | 7 | 0.212 | 0.2 | 2020 | This study |
| France | Etang de Diane | **COR** | 42,13 | 9,54 | 10 | 0.225 | 0.2 | 2020 | This study |
| France | Etang de Diane | *CORS* | 42,71 | 9,54 | 7 | 0.189 | 18.9 | Unknown | Alves Monteiro et al. 2024 |
| Italy | Sardegna | **SARD** | 39,90 | 8,50 | 18 | 0.227 | 0.2 | 2019 | This study |
| Italy | Golfo di Oristano | *ORIS* | 39,54 | 8,29 | 13 | 0.181 | 22.8 | 2021 | Alves Monteiro et al. 2024 |
| Croatia | Zecevo | *ZECE* | 43,67 | 15,87 | 20 | 0.180 | 23.7 | 2017 | Alves Monteiro et al. 2024 |

### Merging and cleaning the datasets

In order to include our results in the European framework, we considered the integration of our new GP with a sequencing dataset previously available at the European level, from which genotyping information could be retrieved from the sequencing data. Therefore, we downloaded the low coverage sequences of 15 populations from Alves Monteiro et al. (2024), namely AGAB, BUNN, CORS, GREV, HAUG, HYPP, LOGS, ORIS, OSTR, PONT, RYAN, TRAL, TOLL, WADD, and ZECE, for a total number of 260 oyster samples (Table 1). These samples are referred to as Sequenced Populations (SP).

We filtered the raw sequences with Fastp (Chen et al. 2018) to remove adapter sequences and polyA and polyG tails from reads, and keep only reads with an average quality of at least 28 (--average_qual 28) and a minimum length of 50 (--length_required 50). We used BWA2 (Vasimuddin et al., 2019) to map the reads onto the flat oyster reference genome (Gundappa et al. 2022), using default parameters. We used SAMBAMBA (Tarasov et al., 2015) to keep only properly paired reads and sort them. We used ANGSD (Korneliussen et al., 2014) to perform genotype calling, using a more relaxed approach than that of Alves Monteiro et al. (2024) in order to retrieve the maximum number of variant sites. Genotypes were called with the following parameters: -doDepth 1 -GL 1 -setMinDepthInd 1 -setMinDepth 30 -setMaxDepth 1200 -SNP_pval 1e-2 -doMaf 1 -doMajorMinor 1 -minMaf 0.05 -doBcf 1 - doPost 1 -doCounts 1 -doGeno 8.

We used Geneious Prime V.2025.0.3 (Drummond et al. 2009) to map the sequences of the loci from the *O. edulis* SNP array to the same reference genome as SP (Gundappa et al. 2022), and used the mapping position to derive the position of the variants of interest (starting position +35bp). We then used BCFtools (Danecek et al. 2021) to filter the BCF and only retain the variable sites that overlapped with the SNPs included in the array. We then combined our own dataset with the filtered BCF file. Considering the important impact of the large inversions present in the European flat oyster genome (Alves Monteiro et al. 2024; Colston-Nepali 2025), we used Plink v1.9 (Purcell et al. 2007) to calculate pairwise LD and remove SNPs with an r^2^ higher than 0.1.

In order to limit bias in the allelic frequency estimations due to the low coverage sequencing dataset (SP), we decided to keep SNPs for which the difference between observed heterozygosity estimates (*Ho*) of GP and SP were not significantly different from zero (p value > 0.05). These estimates were obtained using the R package hierfstat following Nei (1987) equations, and a Kruskal-Wallis rank sum test was applied with the R package stats (function kruskal.test, Hollander and Wolfe 1973), with a significance level set at p=0.05.

### Genetic diversity, relatedness and structure

To assess the overall genetic similarity among individuals, we calculated identity by state (IBS) for all pairs of individuals within each population, using the R package SNPRelate v1.44.0 (Zheng et al. 2012) and the snpgdsIBS function. Intra-population IBS values were then plotted using ggplot2 v4.0.2 (Wickham 2016).

We used the program STRUCTURE (v.2.3.2; Pritchard et al. 2000), based on a Bayesian algorithm that clusters individuals to minimize deviations from Hardy–Weinberg equilibrium. We ran the program 10 times for each number of clusters (*k*) between 1 and 10 using prior population information, with a correlated allele frequency model, and a burn-in of 50,000 iterations followed by 500,000 iterations. Several *k*-estimation methods were used, including the Pr[X|K] method (Pritchard et al. 2000), the ΔK method (Evanno et al. 2005), and the parsimony method thanks to the KFinder program (Wang 2019).

We performed a clustering analysis with a discriminant analysis of principal components (DAPC) using the R package adegenet (Jombart et al., 2010). This procedure consists of running successive k-means with an increasing number of clusters (*k*), after transforming data using a principal component analysis (PCA) performed with the R package ade4 (function dudi.pca). PCA analyses were performed by replacing the missing data by the mean of the allelic frequencies in the populations. A statistical measure of goodness of fit is computed to determine optimal *k*. The samples were assigned to those *k* clusters, which was visualized with a pie chart created using R packages rnaturalearth v1.2.0 (Massicote & South 2026), scatterpie v0.2.6 (Yu 2025) and geosphere v1.6.5 (Hijmans 2026). The k clusters were also represented in the PCA figure with a color code derived from Lapègue et al. (2023).

The differentiation between pairs of populations were estimated with StAMPP R package, thanks to the function stamppFst that calculates pairwise *F_ST_* values along with confidence intervals and p-values between populations according to the method proposed by Wright (1949) and updated by Weir and Cockerham (1984). Isolation By Distance (IBD) was tested with a Mantel test (Mantel, 1967) thanks to the mantel.rtest function of the R package ade4. Finally, to test for possible admixture between clusters identified by STRUCTURE, we used Treemix v1.13 (Pickrell & Pritchard 2012) and the threepop function to calculate f_3_ statistics. For example, if f_3_(A;B,C) Z-score is negative, it indicates that cluster A (the focus cluster) results from the admixture of clusters B and C.

### Outlier loci

To better characterize the geographical clusters obtained through Structure and identify loci that contribute to this structuring, we used three methods. First, we used BAYESCAN (Foll and Gaggioti 2008), a Bayesian approach that uses a logistic regression model to estimate the prior probability that a given locus is under selection. The parameters were: -burn-in 50000; -thinning interval 10; -sample size 750; -resulting total number of iterations 57500; -number of pilot runs 20; -length of each pilot run 5000. Secondly, we performed a DAPC analysis with the two groups of populations and identified the SNPs whose contribution to the axis is greater than the 99th percentile (i.e., the top 1%). Finally, the OutFLANK method of Whitlock and Lotterhos (2015) was applied, using the gl.outflank function of the R package dartR with the default parameters. Any locus identified by at least one method was considered as an outlier. We used the published genome annotation (Gundappa et al. 2022) to functionally characterize the outliers. An outlier was considered as putatively impacting a gene when found within 5kb upstream or downstream of this gene.

## Results

### Handling the bias of the merged dataset

Merging the two datasets and the different filtering steps resulted in a dataset of 2129 SNPs for 747 samples from 35 populations, 487 from GP and 260 from SP, referred to as the Merged Dataset I (Lapègue et al. 2026). We observed a bias in this dataset between GP and SP for the observed heterozygosity estimate *Ho* (Figure 1A) and decided to keep SNPs for which the difference between the heterozygosity estimates of GP and SP was not significantly different from zero (p value > 0.05). From the initial 2129 SNPs in the Merged Dataset I, we ended with 1001 SNPs for the 747 samples (Figure 1B, Table 1) for which we still observe the same tendency, that is however not significant. This reduced dataset, referred as Merged Dataset II, was used for the analyses made at the European level. The percentage of missing data (Fig. S1, Table 1) was higher for the SP (17.6%) than the GP (0.6%).

**Fig. 1.**
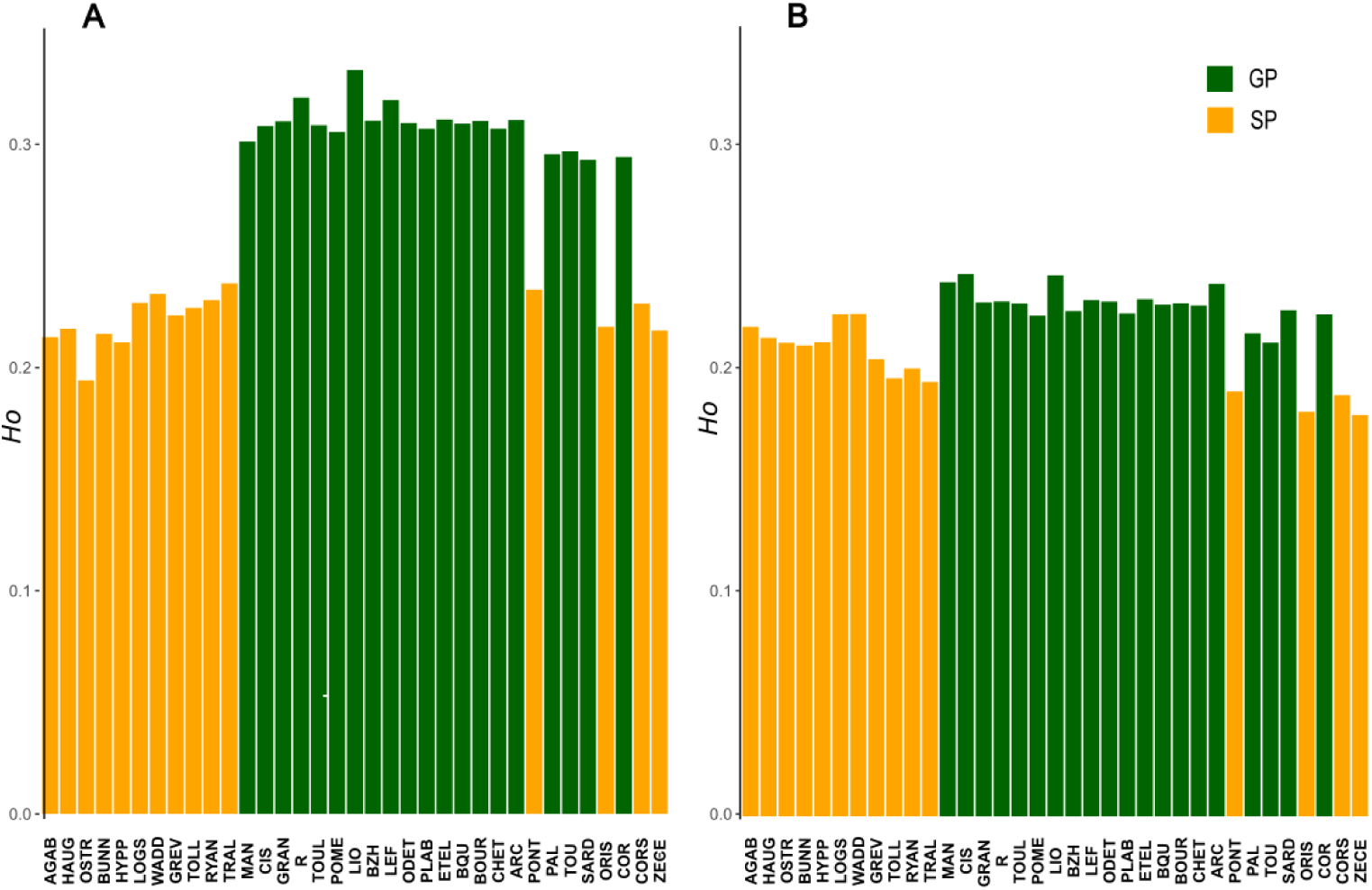
Observed Heterozygosity *Ho* for Merged Dataset I with 2129 SNPs (A) and Merged Dataset II with 1001 SNPs (B). The Genotyped Populations (GP) are colored in green and the Sequenced Populations (SP) populations are colored in orange. The populations are in geographic order, from north to the south and then east, along the coasts.

### The flat oyster clusters in the European context

The clustering analysis with DAPC gave a minimum value of BIC (Bayesian Information Criterion) for *k*=4 but with very close values for *k*=3 and *k*=5 (Fig. S2). In parallel, the STRUCTURE analysis gave *k*=6 as the best *k* estimated with the three methods tested (Table S1). However, considering that *k*=5 also demonstrated strong likelihood (Δ_Parsimony_=0.0097 between *k=5* and *k=6*), we examined all four possibilities (Fig. S3).

For *k*=4, 5 and 6 (Fig. S3 B-C-D), we find again the three clusters that were detected previously by Lapègue et al. (2023): Med_West (former MedW, light blue) present along the coast in the western part of the Mediterranean, ATL (former Atl, dark blue) present along the coast of the French Atlantic and the British Isles, and NS_South (former NS, in red) present along the coast of the North Sea and the British Isles. This renaming of the former NS in NS_South is due to the detection of a NS_North cluster, here in dark red, corresponding to several northern Norwegian populations, also observed in Alves Monteiro (2024). Although being identified as the second-best value by the ΔK method, *k*=3 did not catch the NS_South cluster (in light red), creating a gradient of admixture between the NS_North (in dark red) and ATL (in dark blue) clusters, for HAUG, BUNN, HYPP, LOGS, WADD and GREV populations (Fig. S3A). In the same way, *k*=6 creates the ATL_South cluster (in blue) that shows a gradient of admixture with ATL (Fig. S3C). Moreover, a new cluster was detected, called CHAN_East in pink, corresponding to two very close populations sampled along the Northern French coasts, MAN and CIS (Fig. S3C, and S3D), at the eastern part of the English Channel. Because the clusters appear geographically best defined for *k*=5 (rather than *k*=6 which evidence a sort of latitudinal gradient within a patch, likely due to introgression tails), we chose to represent the different further analyses with this clustering and the corresponding colors (Fig. 2). Furthermore, DAPC analysis indicates that the vast majority of the oysters sampled can be assigned to more than 50% to one of the 5 clusters detected.

**Fig. 2.**
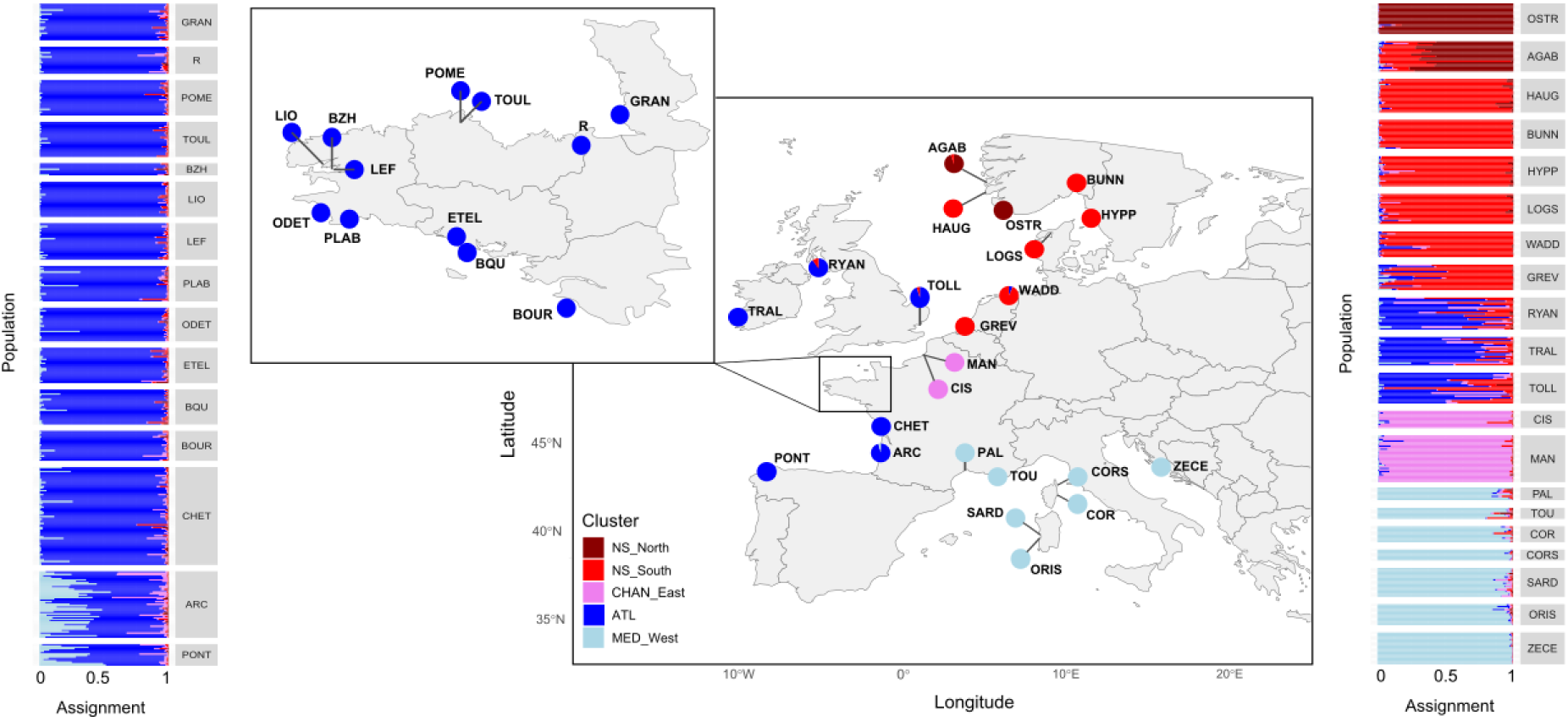
Population genetic structure of flat oyster samples across Europe with *k=5.* Central map indicates sampling locations, and pie charts correspond to the cluster membership of individuals (determined using DAPC analyses). Barplot panels on either side correspond to the individual cluster assignments (determined using STRUCTURE analyses).

### A high homogeneity in Brittany

The PCA showed a clear geographic pattern in the two first axes (Fig. 3A). This geographic pattern was confirmed by the IBD test, demonstrating a positive association between genetic and geographic distance (r=0.58; p = 0.0001). IBD test focusing on ATL populations from Britany to the Pertuis Charentais (from GRAN to CHET), also demonstrated a strong pattern of isolation by distance (r=0.61; p=0.01). This signal likely explains the emergence of an additional ATL cluster (ATL_South) on the STRUCTURE plot for *k*=6, which follows a latitudinal gradient (Fig. S3D).

**Fig. 3.**
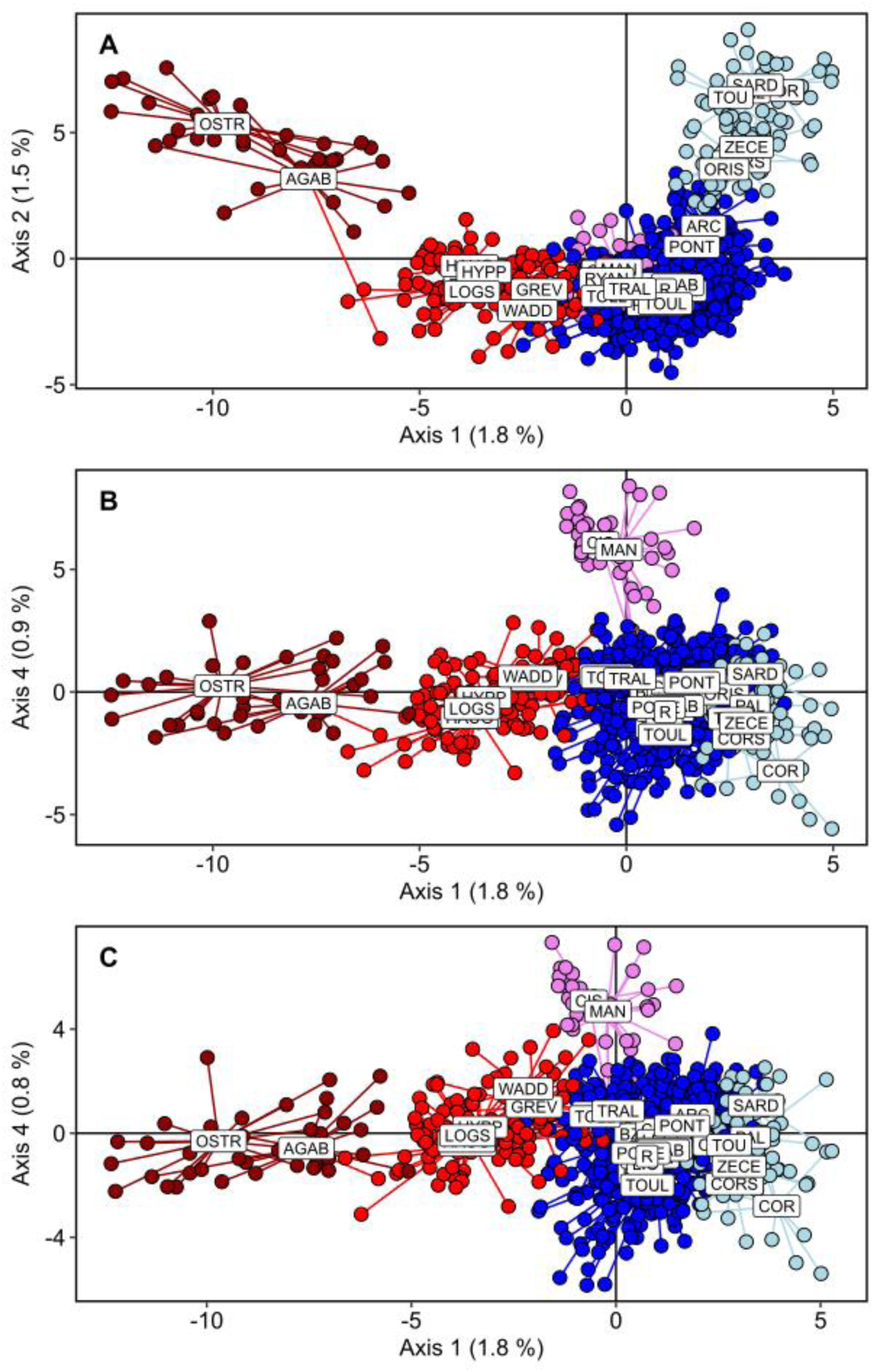
PCA for *k*=5 for A) axes 1 and 2, B) axes 1 and 4, and C) axes 1 and 4 with outliers removed. Colors correspond to the 5 clusters described in Figure 2.

The *F_ST_* matrix (Fig. 4) confirmed that populations from GRAN to CHET have strong genetic similarity with very low *F_ST_* values that were not significantly different from zero for all pairs of populations. The three populations from the British Isles are clearly admixed between the NS_South and ATL clusters (Fig. 2) and in an intermediate position between the NS_South and ATL clusters on the first axis of the PCA (Fig. 3A). Despite belonging to the ATL cluster, ARC and PONT are more differentiated from the other ATL populations (Fig. 4). They appear admixed between the ATL and MED_West clusters (Fig. 2) and in an intermediate position between the NS_South and ATL clusters on the second axis of the PCA (Fig. 3A).

**Fig. 4.**
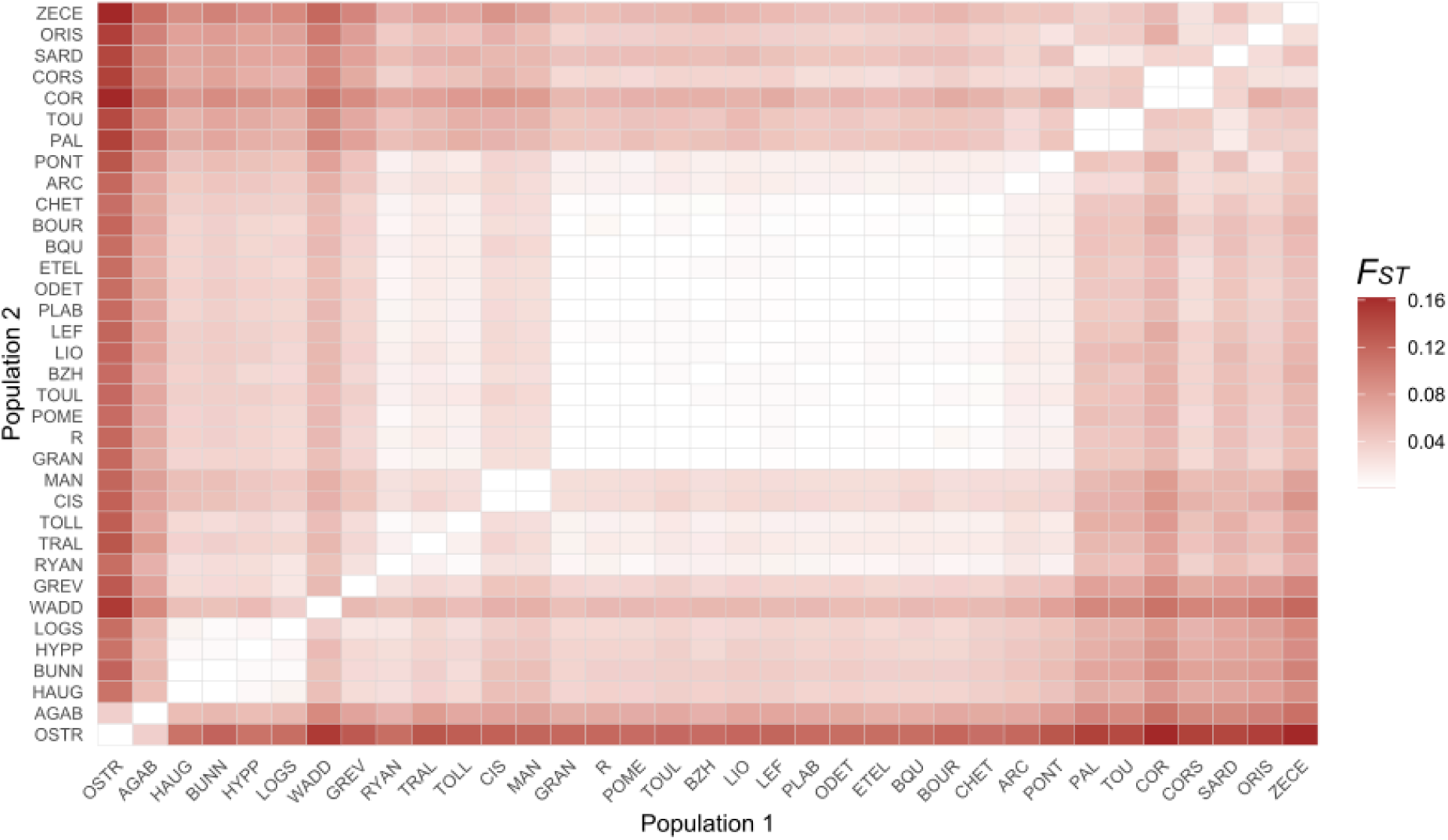
Matrix of pairwise *F_ST_* for all populations (organized geographically). Cells are colored in a gradient corresponding to *F_ST_* values. White cells correspond to *F_ST_* values that are not significantly different from zero.

### A new highly localized cryptic genetic cluster restricted to Northern France

Axis 4 on the PC clearly separated the two populations of CHAN_East (in pink) from all the others (Fig. 3B). They form a separate cluster in the STRUCTURE analysis for *k*=5 (Fig. 2) and are significantly differentiated from the other ATL populations (Fig. 4). While PCA analyses were performed by replacing the missing data by the mean of the allelic frequencies in the populations, we explored a potential impact of the differences (percentage of missing data, *Ho*) between GP and SP. This potential impact was explored using samples from the western Mediterranean Sea: PAL, TOU, COR, SARD (GP), and CORS, ORIS and ZECE (SP). They appear to show separation in two groups on the second axis of the PCA (Fig. 3A). Those two groups of Mediterranean populations are characterized by high/low levels of missing data for GP/SP, and low/high heterozygosity for GP/SP (Fig. S1; Fig. S4 A and B), which appears to slightly influence the position of the individuals on the PCA. For *Ho*, the MANOVA model was significant for the first 4 axes (pvalue=1.1 x 10^-7^), the principal amount of variance explained being on axis 2 (3%). For the percentage of missing data, the model was also significant the first 4 axes (pvalue < 1.1 x 10^-16^), the principal amount of variance explained being on axis 1 (11.5%). However, COR and CORS, from GP and SP respectively, were sampled at the same place and demonstrate a pairwise *F_ST_* that is not significantly different from zero (Fig. 4). In the STRUCTURE analysis, all Mediterranean populations clearly cluster in MED_West regardless of *k* (Fig. S3). Altogether, the different analyses performed on the Merged Dataset II were able to clearly characterize the MED_West cluster. Such an impact of the high level of missing data and low heterozygosity on PCA does not however concern MAN and CIS (both from GP), in the CHAN_East cluster, both demonstrating a low percentage of missing data and relatively high *Ho* (Fig. S1; Fig. S4 C and D).

Being at the boundary of the ATL and NS_South clusters, we more deeply investigated the characteristics of MAN and CIS in order to better understand the origin of this cluster. Outlier analyses identified 16 outlier loci (Fig. S5) between the MAN+CIS populations (CHAN_East cluster) and all other populations in the dataset (Table S2). These outliers are not physically regrouped in a specific region, and are located on 9 of the 10 chromosomes of the genome assembly. Two loci are located in intergenic areas while 14 located near or within genes. However, even when removing these outliers, the separation of CHAN_East from the other clusters is still clearly visible on the PCA (Fig. 3C), and their removal only decreased the variance explained by axis 4 from 0.9% to 0.8%.

The population tree inferred using Treemix (Fig. 5) supported a model of one migration event from MED_West towards ATL, with no migration from or to CHAN_East. Furthermore, the branch length was smaller for ATL indicating of a higher admixture for this cluster. This is also visible in the STRUCTURE plot with an introgression of the NS_South and MED_West on the edges of the ATL distribution (Fig. 2). Finally, f3 statistics confirm that CHAN_East cluster does not result from the admixture of ATL with NS_South, the Z-score estimates being all significantly positive (focus cluster: NS_South, Z-score = 6.64; focus cluster: CHAN_East, Z-score = 6.63; focus cluster: ATL, Z-score = 2.07).

**Fig. 5.**
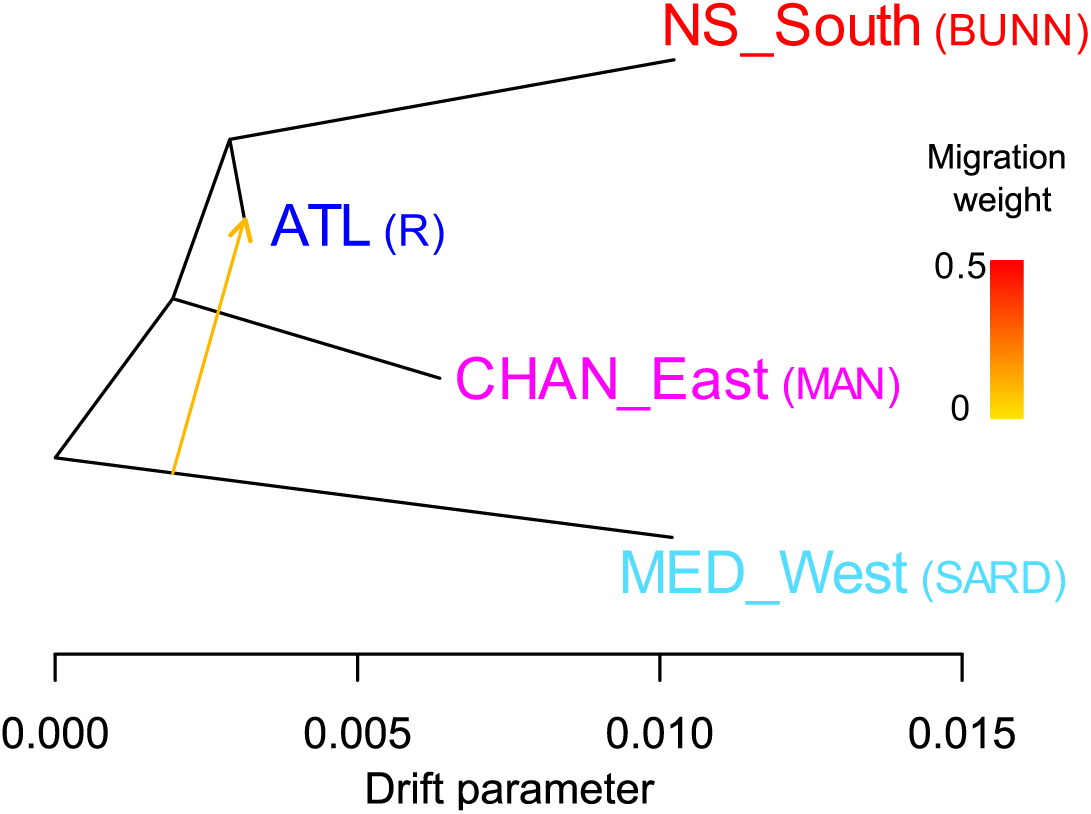
Evolutionary history of four populations representing four genetic clusters using the population graph approach of TREEMIX.

Because IBS is very sensitive to estimations of Ho, and since discrepancies observed in *Ho* between SP and GP (Fig. 1) may still exist in the Merged dataset II, we focused on analysis on the initial SNP Array Dataset, containing 5090 SNPs and including populations from CHAN_East, ATL, and MED_West. While IBS estimates were slightly higher for CHAN_East compared to ATL, they do not markedly differ from MED_West estimates (Fig. 6). This indicates that MED_West and CHAN_East could have similar effective population sizes and did not experience a recent reduction in population size. Those observations, together with the admixture results (Fig. 5) support the conclusion that CHAN_East populations (MAN and CIS) do not follow the same admixture pattern that the other French (ATL) populations and constitute a real separate cluster.

**Fig. 6.**
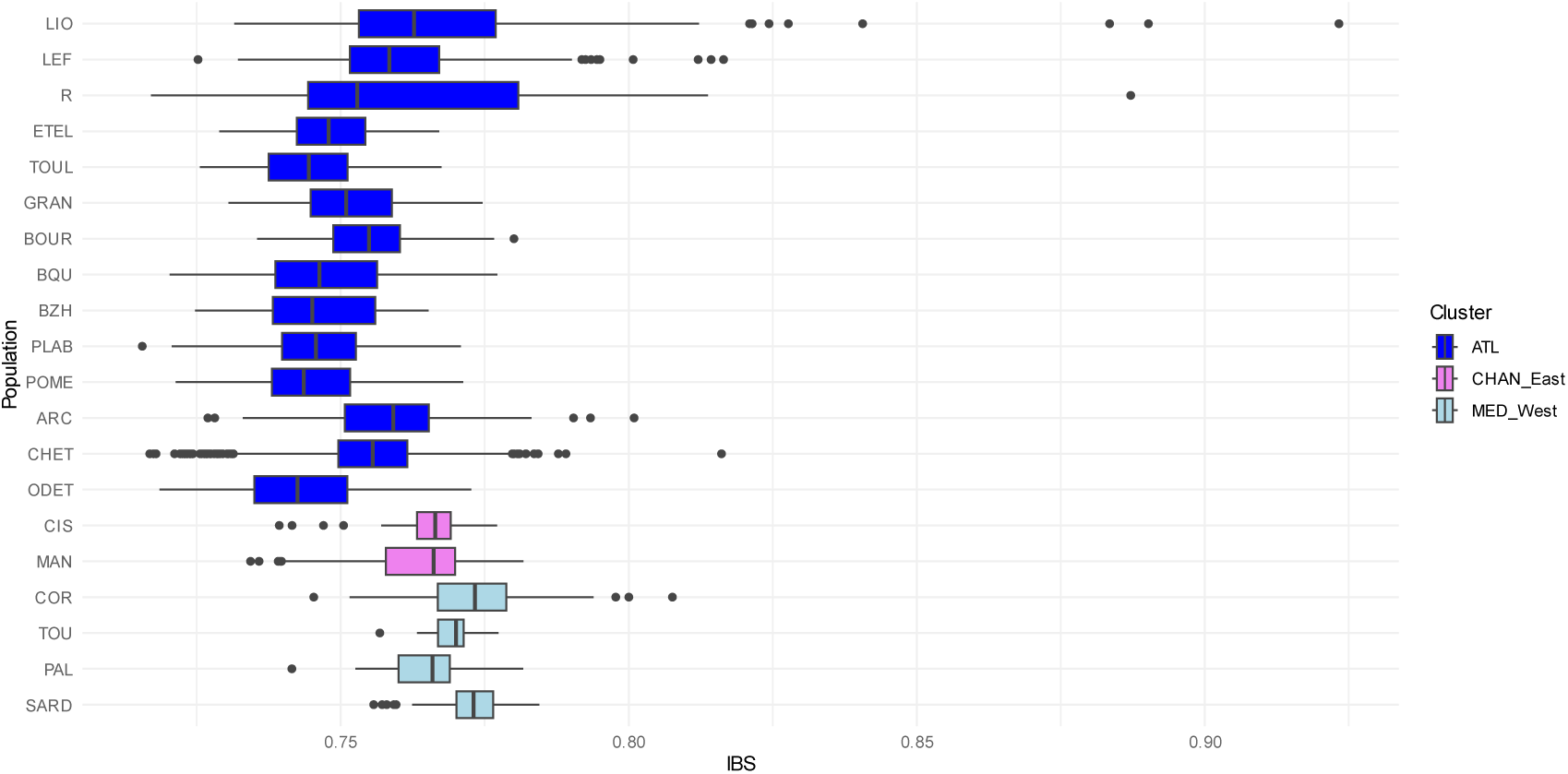
Relatedness (IBS) of the GD populations.

## Discussion

Through the integration of SNP array and low-coverage sequencing datasets, our study proposes a synthetic overview of the population genetic structure of the European flat oyster (*Ostrea edulis*) across Europe. While such integrative approaches present several challenges, they also offer a unique opportunity to consolidate efforts among research groups at the European scale. Most of the previously identified clusters in Western Europe were confirmed. In France, populations assigned to these clusters (ATL and MED_West) exhibit notable within patch homogeneity. However, two key findings emerge: (1) the extensive overlap zone between MED_West and ATL, spanning at least from southern Portugal to southern France, and (2) the detection of a novel, clearly distinct cryptic cluster between NS_South and ATL, whose geographic range remains to be better delineated. These insights are critical for informing management decisions, particularly as restoration and conservation plans are currently being implemented.

### 1) Integrating diverse genomic datasets: challenges, biases, and opportunities for population genetic analyses

A range of genetic and genomic tools are now available for studying European flat oyster (*Ostrea edulis*) populations. Following the use of allozymes and microsatellites, population geneticists have adopted Illumina GoldenGate® and Affymetrix Axiom® technologies to develop and apply SNP arrays for this species (Lapègue et al. 2013; Gutierrez et al. 2017; Vera et al. 2019; Lapègue et al. 2023; Valk et al. 2026b), as well as low-coverage whole-genome sequencing (lcWGS) (Alves Monteiro et al. 2024; Robert et al. 2025). Although these tools have been applied to different samples at regional or European scales, they have collectively revealed a coherent broad-scale genetic structure across Europe. Our study builds on this foundation by utilizing the medium-density combined-species Affymetrix Axiom SNP array (Gutierrez et al. 2017), which includes 14,950 SNPs specific to *O. edulis*. After applying stringent filtering criteria, we retained 5,090 polymorphic SNPs for population genetics analyses. Beyond merely comparing our results with those of previous studies, we aimed to integrate our SNP array data with lcWGS datasets, particularly the dataset from Alves Monteiro et al. (2024), which includes northern European clusters not represented in our sampling. However, integrating these two data types presents challenges. SNPs called from lcWGS data may be less accurate, particularly for rare alleles and in a species characterized by widespread polymorphic duplications that generate numerous paralogous variants (Colston-Nepali et al. 2025). In contrast, Affymetrix Axiom arrays provide highly reproducible genotypes and are largely depleted of paralogous loci owing to probe design and quality-control procedures, albeit at the cost of more limited genomic coverage. Thus, a key consideration is determining the optimal number and quality of SNPs to include in a combined analysis.

To address the issue of SNP quantity, we initially opted for a more lenient genotype-calling approach than that used by Alves Monteiro et al. (2024) to maximize the number of variant sites retrieved. This is generally not a recommended practice, and Colston-Nepali et al. (2025) have in fact advocated for systematic filtering of whole-genome sequencing data using NGSparalog (Linderoth 2018). However, we here rely on SNP array data to select SNPs. We furthermore excluded groups of SNPs in strong linkage disequilibrium located in known chromosomal rearrangements in this species (Sambade et al. 2022; Lapègue et al. 2023; Alves Monteiro et al. 2024; Ewers et al. 2025; Colston-Nepali 2025). This resulted in 2029 shared SNPs, a relatively high number given the low initial number from the SNP array (5090) and the low coverage of the sequencing dataset. However, since not all SNPs could be called across all SP individuals, the missing data rate was higher in SP than in GP. Additionally, we identified a bias in Merged Dataset I between GP and SP for the observed heterozygosity (*Ho*) estimate, which persisted at a much lower intensity in Merged Dataset II. Issues relating to bias estimates of heterozygosity and a high rate of missing data is well known for lcWGS especially when compared to SNP array datasets (Duntsch et al 2021; Kardos and Waples 2024), and it is nowadays recommended to avoid hard calling of genotypes from lcWGS, using instead the genotype likelihood (Lou et al, 2021; Kardos and Waples 2024). However, this is not possible when analyzing lcWGS and SNP array together, and only additional filtering of loci can allow for the integration of both datasets.

The availability of replicated populations in very close geographical proximity in the Western Mediterranean Sea (if not the same samples used as true biological replicates) enabled us to characterize the bias observed in the PCA and assess its impact on the observed genetic patterns. This allowed us to definitively confirm the presence of the new CHAN_East cluster and proceed with our analysis by focusing solely on the SNP array dataset, now including this newly identified cluster. Nevertheless, this bias must be acknowledged as a persistent challenge when integrating these two types of data, and it may complicate future efforts to combine them (although there have been recent developments in genotype imputation from lcWGS likelihoods, for certain applications; Triay et al 2025). Despite this, such integration remains highly valuable for improving our understanding of the genetic structure of French populations within the broader European range of the species. To advance this goal, scientific collaborations should aim to pool their genomic tools and some of their samples, which can serve as references, for more accurate and comprehensive genetic pattern analyses.

### 2) A high homogeneity within the different French clusters

Our study enhances the understanding of the genetic landscape of European flat oyster populations by significantly expanding the number of analyzed populations in France, particularly in Brittany, where the species holds emblematic status. At the national level, *O. edulis* has been documented in Normandy, Nouvelle-Aquitaine, and several Mediterranean lagoons, but its highest abundance is observed in Brittany (Hussenot et al. 2014; Duchêne et al. 2015). Even in this region, however, observations are predominantly limited to isolated individuals, typically at stage 0 (rolling) or stage 1 (fixed but at very low densities). Occasionally, small, highly isolated colonies (stage 2) are encountered, and only exceptionally are aggregations of significant density (stage 3) found. No population has been classified as stage 4, which would correspond to the presumed original reef habitat (Pouvreau et al. 2023; zu Ermgassen et al. 2021). Given the rarity of these animals, sampling posed a challenge and was conducted in strict adherence to the 3Rs principles (Replacement, Reduction, Refinement) in scientific research. The number of specimens collected ranged from 7 up to 29 with the exception of ARC and CHET (72 and 49 respectively) for which temporal samples were pooled. The same individuals were utilized for biometric, pathological, and genetic analyses within the FOREVER and REFONA project (Pouvreau et al. 2021; Carpentier et al. 2024). From Normandy (GRAN) to Nouvelle-Aquitaine (CHET), 13 populations exhibited high genetic similarity within the previously identified ATL cluster (Lapègue et al. 2023; Alves Monteiro et al. 2024), and follow an isolation-by-distance model. Despite historical stock transfers between the Bay of Quiberon and the Bay of Brest to support shellfish farming, the genetic structure of the ATL cluster appears to have remained largely stable over the past centuries. Furthermore, this cluster exhibits greater admixture compared to the CHAN_East and MED_West clusters, likely reflecting an ancient evolutionary history (Lapègue et al. 2023).

The Mediterranean French populations were grouped with the other Mediterranean populations within the MED_West cluster, which also exhibits high genetic homogeneity. Although our analysis included reference populations from this region (Alves Monteiro et al. 2024), we did not observe the separation between their Cluster 1 (Adriatic Sea, including Croatian sampling sites) and Cluster 2 (Mediterranean Sea, including French-Corsican and Italian-Sardinian sampling sites). As previously discussed, this discrepancy may stem from differences in the types of data merged, but it is more likely to be due to the strong reduction in total number of loci analyzed here. Indeed, in our study, the CORS, ZECE, and ORIS populations from Alves Monteiro et al. (2024) were clearly grouped together, despite being sampled along the Corsican, Adriatic, and Sardinian coasts, respectively. This suggests that the limited number of markers in our study (1,001 SNPs) were insufficient to detect the two distinct clusters identified by Alves Monteiro et al. (2024), which were based on over 1.4 million polymorphic sites. Once again, this underscores the value of including reference populations in such analyses. While historical data suggest that the species was unevenly distributed, except possibly in the northern Adriatic (Thurstan et al. 2024), future sampling efforts using a larger number of markers should enable a more refined analysis of the genetic structure in the western Mediterranean.

### 3) The case of admixed populations

The three British Isles populations (RYAN, TRALL, and TOLL) belong to the ATL cluster but exhibit admixture with the NS_South cluster, confirming the observations of Alves Monteiro et al. (2024). This admixed pattern suggests that these populations are close to a genetic boundary between the two clusters. In the southern Bay of Biscay, we also observed introgression in Arcachon Bay (ARC) and Pontedeume (PONT), located in Nouvelle-Aquitaine (southern French Atlantic coast) and Galicia (Spain), respectively. This suggests that the genetic boundary previously identified between the ATL and MED_West clusters may be more extensive than initially thought. Hence, Portuguese samples have already shown signals of introgression, indicating gene flow between these two clusters (Launey et al. 2002; Lapègue et al. 2023). These samples were more closely related to Western Mediterranean populations than to Atlantic ones. In Galicia, PONT serves as a reference sample, as it was originally part of a study that included five other Galician samples (Vera et al. 2016), where it was designated as SP-PNTD (later renamed PONT in Alves Monteiro et al., 2024). In Vera et al. (2016), Galician samples formed a distinct cluster, separated from another corresponding to the ATL cluster in our analysis. Furthermore, when Mediterranean samples are added (Alves Monteiro et al. 2024), PONT and another Galician sample (RIAE) exhibited the same admixed pattern between ATL and MED_West that we observe here. Together, these findings suggest that the entire Galician region may be introgressed by MED ancestry. Surprisingly, ARC displayed the same admixed pattern as PONT, extending this admixture zone as far north as southern France. The next northern population sampled (CHET), located more than 100 km further north, showed no admixture and was clearly assigned to the ATL cluster. Based on these observations, we can now conclude that the overlap zone between MED_West and ATL extends at least from southern Portugal to southern France.

### 4) A new highly localized cryptic genetic cluster in Northern France

Eight historical beds of flat oysters have been identified in the Hauts-de-France region, where the MAN and CIS samples are located (Thurstan et al. 2024). However, this number is relatively low compared to the 109 beds recorded in the early 1900s along the entire French coastline. This area had not been well studied until now, primarily due to the rarity of these oysters. Nevertheless, these two samples indicate at least a residual presence of one of these beds. Although geographically close, MAN corresponds to oysters dredged at a depth of approximately 60 meters, while CIS corresponds to oysters collected by hand during low tide. Both, however, belong to the newly identified CHAN_East cluster. Our genetic analyses show that this new cluster does not represent an admixture zone between the ATL and NS_South clusters, as observed in the British Isles. Its origin remains somewhat enigmatic. It could correspond to a bed considered extinct, such as the one rediscovered through museum samples in the Wadden Sea (Ewers et al. 2025), with MAN belonging to the bed itself and CIS representing an intertidal extension of it. The outlier markers identified account for only a small portion of the variation along the PCA axis that characterizes the separation between this cluster and other populations. However, among the 14 (out of 16) outliers located in genes, trichohyalin-like was recently identified in the Pacific purple sea urchin as one of the outliers differentiating intertidal and subtidal habitats (Rumberger et al. 2025). In humans, this protein is primarily found in hair follicles, which are specialized structures in the skin where hair growth occurs. It may be of interest to further characterize the function of such genes in marine invertebrates.

### 5) Conservation and restoration issues

Several distinct genetic clusters of the European flat oyster (*Ostrea edulis*) have been identified and characterized across Western Europe, meeting criteria for Evolutionary Significant Units (ESUs) (Ryder 1986). Intraspecific genetic diversity is known to enhance adaptive capacity (DeWoody et al. 2021), yet genetic criteria are not explicitly included in OSPAR or IUCN assessments. Recent studies propose incorporating ESUs into IUCN frameworks to better protect genetic diversity and improve conservation management (Norderhaug et al. 2024; van Oosterhout et al. 2025). While current studies, including ours, primarily analyze selectively neutral molecular markers, they do not capture adaptive or quantitative genetic variation (Camara and Vadopalas 2009). Testing for local adaptation remains challenging due to difficulties in measuring phenotypic traits, particularly fitness. A conservative approach would treat genetic clusters as locally adapted units, but populations within clusters may still exhibit local adaptation for quantitative traits even when gene flow is detected at neutral markers. Increasing genomic coverage may help, but phenotypic data and common garden experiments are essential to complement genomic studies.

In Brittany, where oyster beds persist, restoration focuses on facilitating settlement. In Nouvelle-Aquitaine, recent studies reveal highly fragmented, relict populations and the absence of functional reef habitats (Carpentier et al. 2026), necessitating targeted conservation strategies. Restoration efforts in Pertuis Charentais aim to reinforce populations and improve habitat connectivity, while in Arcachon Bay, preserving the Western Mediterranean genetic introgression might be considered (Carpentier et al. 2026). Current actions rely on pre-seeded substrates from local broodstock to maintain genetic integrity while addressing ecological constraints like substrate scarcity and low reproductive density. The risk of outbreeding depression highlights the need to prioritize local broodstock for supplementation. However, the use of hatchery-produced oysters may increase kinship and inbreeding (Lallias et al. 2010), as seen in Dutch populations (Valk et al. 2026b). Along the French Mediterranean, restoration programs are lacking, and further sampling is needed to clarify genetic structure, especially given the coexistence with *O. stentina* (González-Wangüemert et al. 2004; Lapègue et al. 2006). In Hauts-de-France, a newly identified cluster with clear genetic differentiation requires range assessment, particularly as Belgium’s restoration efforts consider North Sea broodstocks.

Now is the time to assess *O. edulis* reef habitats across France, following Brittany and Nouvelle-Aquitaine’s lead (Pouvreau et al. 2021; Carpentier et al. 2026). This aligns with the EU Nature Restoration Regulation (NRR; European Union 2024), which mandates ambitious national restoration plans. Minimum viable habitat thresholds are being estimated for *O. edulis* reefs in the Atlantic, English Channel, North Sea, and Mediterranean (Laforge et al. 2026). The genetic characterization provided by this study will support evidence-based decision-making for conservation and restoration.

## Author Contributions

All authors contributed to the study conception and design. Material preparation and data collection were performed by Sylvie Lapègue, Florence Cornette, Serge Heurtebise, Stéphane Pouvreau and Cynthia Carpentier. Analyses were performed by Sylvie Lapègue, Lila Colston-Népali, Nicolas Bierne and Céline Reisser. The first draft of the manuscript was written by Sylvie Lapègue and all authors commented on previous versions of the manuscript. All authors read and approved the final manuscript.

## Data Availability

The datasets generated during and/or analyzed during the current study are available in the SEANOE (sea scientific open data publication) repository (Lapègue et al. 2026; https://doi.org/10.17882/117262).

## Acknowledgements

This work was supported by FOREVER and REFONA projects. We would like to thank Josiane Popovsky, Guillaume Ortega, Gaël Oudot, and Johan Vieira from CAPENA (Centre pour l’Aquaculture, la Pêche et l’ENvironnement de Nouvelle-Aquitaine), divers from the Arcachon Bay Marine Nature Park, Céline Rolet from GEMEL (Groupe d’Etude des Milieux Estuariens et Littoraux), Ms. Baudéan from CRPMEM (Comité Régional des Pêches Maritimes de Normandie), and M. Ulrick Comtesse for collecting the oyster samples. We would also like to thank Laëtitia Bernard and Auriane Lecler for their participation in the sampling and DNA extraction. The authors acknowledge the Pôle de Calcul et de Données Marines (PCDM; https://wwz.ifremer.fr/en/Research-Technology/Research-Infrastructures/Digital-infrastructures/Computation-Centre) for providing DATARMOR computing and storage resources.

## Notes

https://doi.org/10.17882/117262

### Competing Interest Statement

The authors have declared no competing interest.

## Reference list

Alves Monteiro HJ, Bekkevold D, Pacheco G, et al (2024) Genome-Wide Population Structure in a Marine Keystone Species, the European Flat Oyster (*Ostrea edulis*). Molecular Ecology 34:e17573. 10.1111/mec.17573

Beck MW, Brumbaugh RD, Airoldi L, et al (2011) Oyster Reefs at Risk and Recommendations for Conservation, Restoration, and Management. BioScience 61:107–116. 10.1525/bio.2011.61.2.5

Bersoza Hernández A, Brumbaugh RD, Frederick P, et al (2018) Restoring the eastern oyster: how much progress has been made in 53 years? Frontiers in Ecology and the Environment 16:463–471. 10.1002/fee.1935

Blanc F, Jaziri H, Durand P (1986) Isolement génétique et taxonomie des huîtres plates dans une lagune du sud de la Méditerranée occidentale. Comptes Rendu de l’Académie des Sciences de Paris, Série III 303:207–210.

Botta R, Asche F, Borsum JS, Camp EV (2020) A review of global oyster aquaculture production and consumption. Marine Policy 117:103952. 10.1016/j.marpol.2020.103952

Bromley C, McGonigle C, Ashton EC, Roberts D (2016) Bad moves: Pros and cons of moving oysters – A case study of global translocations of *Ostrea edulis* Linnaeus, 1758 (Mollusca: Bivalvia). Ocean & Coastal Management 122:103–115. 10.1016/j.ocecoaman.2015.12.012

Camara MD, Vadopalas B (2009) Genetic Aspects of Restoring Olympia Oysters and Other Native Bivalves: Balancing the Need for Action, Good Intentions, and the Risks of Making Things Worse. Journal of Shellfish Research 28:121–145. 10.2983/035.028.0104

Carpentier C, Vieira J, Bernard L, Lecler A, Barbier P, Arzul I, Oudot G, Bodin P, Weiller Y, Leleu K. 2024. Inventaire et caractérisation des populations résiduelles d’huîtres plates en Nouvelle-Aquitaine. Projet REFONA – Restauration & Conservation de l’huître plate en Nouvelle-Aquitaine. Rapport d’étude CAPENA. 97 pp

Carpentier C, Vieira J, Barbier P, Oudot G, Fauvel T, Leleu K, Weiller Y, Dupuy C, Arzul I, Pouvreau S (2026). Rediscovering Ostrea edulis in Nouvelle-Aquitaine (France): From historical records to contemporary distribution patterns. Submitted

Chen S, Zhou Y, Chen Y, Gu J (2018) fastp: an ultra-fast all-in-one FASTQ preprocessor. Bioinformatics 34:i884–i890. 10.1093/bioinformatics/bty560

Coen LD, Luckenbach MW (2000) Developing success criteria and goals for evaluating oyster reef restoration: Ecological function or resource exploitation? Ecological Engineering 15:323–343. 10.1016/S0925-8574(00)00084-7

Cole VJ, Harasti D, Lines R, Stat M (2022) Estuarine fishes associated with intertidal oyster reefs characterized using environmental DNA and baited remote underwater video. Environmental DNA 4:50–62. 10.1002/edn3.190

Colsoul B, Boudry P, Pérez-Parallé ML, et al (2021) Sustainable large-scale production of European flat oyster (*Ostrea edulis*) seed for ecological restoration and aquaculture: a review. Reviews in Aquaculture 13:1423–1468. 10.1111/raq.12529

Colston-Nepali L (2025) Variations structurales et héterogénéité des paysages génomiques de différenciation parallèle chez l’huître plate européenne (*Ostrea edulis*). Structural variants and a heterogeneous genomic landscape of parallel differentiation in the European flat oyster (Ostrea edulis). Dissertation. University of Montpellier

Colston-Nepali L, Lapègue S, Bierne N (2025) *k*-mer spectra and allelic coverage analyses reveal pervasive polymorphic duplications in *Ostrea edulis*. BioRxiv. 10.1101/2025.06.24.661118

Danecek P, Bonfield JK, Liddle J, et al (2021) Twelve years of SAMtools and BCFtools. Gigascience 10:giab008. 10.1093/gigascience/giab008

DeWoody JA, Harder AM, Mathur S, Willoughby JR (2021) The long-standing significance of genetic diversity in conservation. Molecular Ecology 30:4147–4154. 10.1111/mec.16051

Diaz-Almela E, Boudry P, Launey S, et al (2004) Reduced Female Gene Flow in the European Flat Oyster *Ostrea edulis*. Journal of Heredity 95:510–516. 10.1093/jhered/esh073

Drummond AJ, Ashton B, Cheung M, et al (2009). Geneious v4.8. [software]. http://www.geneious.com

Duarte CM, Agusti S, Barbier E, et al (2020) Rebuilding marine life. Nature 580:39–51. 10.1038/s41586-020-2146-7

Duchêne J, Bernard I, Pouvreau S (2015) Vers un retour de l’huitre indigène en rade de Brest. Espèces 16:51–57. https://archimer.ifremer.fr/doc/00270/38085/

Duntsch L, Whibley A, Brekke P, et al (2021) Genomic data of different resolutions reveal consistent inbreeding estimates but contrasting homozygosity landscapes for the threatened Aotearoa New Zealand hihi. Molecular Ecology 30:6006–6020. 10.1111/mec.16068

European Union (2024) Regulation (EU) 2024/1991 of the European Parliament and of the Council of 27 June 2024 on nature restoration. Official Journal of the European Union. https://eurlex.europa.eu/legal-content/EN/TXT/?uri=CELEX%3A32024R1991. Assessed 27 May 2026

Evanno G, Regnaut S, Goudet J (2005) Detecting the number of clusters of individuals using the software structure: a simulation study. Molecular Ecology 14:2611–2620. 10.1111/j.1365-294X.2005.02553.x

Ewers C, Brandis D, da Silva N, et al (2025) Museomics of an extinct European flat oyster population. Scientific Reports 15:13906. 10.1038/s41598-025-96743-8

Foll M, Gaggiotti O (2008) A Genome-Scan Method to Identify Selected Loci Appropriate for Both Dominant and Codominant Markers: A Bayesian Perspective. Genetics 180:977–993. 10.1534/genetics.108.092221

González-Wangüemert M, Perez-Ruzafa A, Rosique M, Ortiz A (2004) Genetic differentiation in two cryptic species of Ostreidae, *Ostrea edulis* (Linnaeus, 1758) and *Ostreola stentina* (Payraudeau, 1826) in Mar Menor Lagoon, southwestern Mediterranean Sea. Nautilus 118:103–111

Gundappa MK, Peñaloza C, Regan T, et al (2022) Chromosome level reference genome for European flat oyster (*Ostrea edulis* L.). bioRxiv 2022.06.26.497633. 10.1101/2022.06.26.497633

Gutierrez AP, Turner F, Gharbi K, et al (2017) Development of a Medium Density Combined-Species SNP Array for Pacific and European Oysters (*Crassostrea gigas* and *Ostrea edulis*). G3 Genes|Genomes|Genetics 7:2209–2218. 10.1534/g3.117.041780

Hijmans R (2026) geosphere: Spherical Trigonometry. R package version 1.6-7, https://github.com/rspatial/geosphere.

Hollander M, Wolfe DA (1973) Nonparametric Statistical Methods. New York: John Wiley & Sons. Pages 115–120.

Hussenot M, Pouvreau S, Duchêne J, Freulon H, Arzul I, Lapègue S (2014) Synthèse PERLE. Programme d’Expérimentation et de recherche sur L’huître plate Ostrea edulis. https://archimer.ifremer.fr/doc/00249/36060/

Jombart T, Devillard S, Balloux F (2010) Discriminant analysis of principal components: a new method for the analysis of genetically structured populations. BMC Genetics 11:94. 10.1186/1471-2156-11-94

Kamermans P, Anteau F, Didderen K, et al (2025) Novel settlement substrates for European flat oyster (*Ostrea edulis*) restoration. Ecological Engineering 212:107532. 10.1016/j.ecoleng.2025.107532

Kardos M, Waples RS (2025) Low-coverage sequencing and Wahlund effect severely bias estimates of inbreeding, heterozygosity and effective population size in North American wolves. Molecular Ecology 34:e17415. 10.1111/mec.17415

Korneliussen TS, Albrechtsen A, Nielsen R (2014) ANGSD: Analysis of Next Generation Sequencing Data. BMC Bioinformatics 15:356. 10.1186/s12859-014-0356-4

Laforge D, Le Moal M, Aulay M, et al (2026) Estimation préliminaire des Surfaces de Référence Favorable des GTH marins dans le cadre de la préparation du Plan National de Restauration 2027 : méthodes et estimations chiffrées des SRF. PatriNat (OFB-MNHN-CNRS-IRD), Paris, 41 pp

Laikre L, Schwartz MK, Waples RS, Ryman N (2010) Compromising genetic diversity in the wild: unmonitored large-scale release of plants and animals. Trends in Ecology & Evolution 25:520–529. 10.1016/j.tree.2010.06.013

Lallias D, Taris N, Boudry P, et al (2010) Variance in the reproductive success of flat oyster *Ostrea edulis* L. assessed by parentage analyses in natural and experimental conditions. Genetics Research 92:175–187. 10.1017/S0016672310000248

Lapègue S, Ben Salah I, Batista FM, et al (2006) Phylogeographic study of the dwarf oyster, *Ostreola stentina*, from Morocco, Portugal and Tunisia: evidence of a geographic disjunction with the closely related taxa, *Ostrea aupouria* and *Ostreola equestris*. Mar Biol 150:103–110. 10.1007/s00227-006-0333-1

Lapègue S, Reisser C, Harrang E, et al (2023) Genetic parallelism between European flat oyster populations at the edge of their natural range. Evolutionary Applications 16:393–407. 10.1111/eva.13449

Lapègue S, Cornette F, Heurtebise S, Reisser C (2026). SNP datasets for the genetic analysis of European flat oyster populations. SEANOE. 10.17882/117262

Launey S, Ledu C, Boudry P, et al (2002) Geographic Structure in the European Flat Oyster (*Ostrea edulis* L.) as Revealed by Microsatellite Polymorphism. Journal of Heredity 93:331–351. 10.1093/jhered/93.5.331

Lou RN, Jacobs A, Wilder AP, Therkildsen NO (2021) A beginner’s guide to low-coverage whole genome sequencing for population genomics. Molecular Ecology 30:5966–5993. 10.1111/mec.16077

Mantel N (1967) The detection of disease clustering and a generalized regression approach. Cancer Research, 27, 209–220.

Massicotte P, South A (2026). rnaturalearth: World Map Data from Natural Earth. R package version 1.2.0.9000, https://docs.ropensci.org/rnaturalearth/

McAfee D, Connell SD (2021) The global fall and rise of oyster reefs. Frontiers in Ecology and the Environment 19:118–125. 10.1002/fee.2291

Nei M (1987) Molecular Evolutionary Genetics. Columbia University Press. 512 pp

Neild RR (1995) The English, the French, and the oyster. Quiller Press, London, 212 pp

Norderhaug KM, Knutsen H, Filbee-Dexter K, et al (2024) The International Union for Conservation of Nature Red List does not account for intraspecific diversity. ICES Journal of Marine Science 81:815–822. 10.1093/icesjms/fsae039

Pickrell JK, Pritchard JK (2012) Inference of Population Splits and Mixtures from Genome-Wide Allele Frequency Data. PLOS Genetics 8:e1002967. 10.1371/journal.pgen.1002967

Pogoda B, Brown J, Hancock B, et al (2019) The Native Oyster Restoration Alliance (NORA) and the Berlin Oyster Recommendation: bringing back a key ecosystem engineer by developing and supporting best practice in Europe. Aquatic Living Resources 32:13. 10.1051/alr/2019012

Pouvreau S, Cochet H, Fabien A, et al (2021). Inventaire, diagnostic écologique et restauration des principaux bancs d’huitres plates en Bretagne : le projet FOREVER. Rapport final. Contrat FEAMP 17/2215675. 10.13155/79506

Pouvreau S, Lapègue S, Arzul I, Boudry P (2023) Fifty years of research to counter the decline of the European flat oyster (*Ostrea edulis*): a review of French achievements and prospects for the restoration of remaining beds and revival of aquaculture production. Aquat Living Resour 36:13. 10.1051/alr/2023006

Pritchard JK, Stephens M, Donnelly P (2000) Inference of Population Structure Using Multilocus Genotype Data. Genetics 155:945–959. 10.1093/genetics/155.2.945

Purcell S, Neale B, Todd-Brown K, et al (2007) PLINK: a tool set for whole-genome association and population-based linkage analyses. The American Journal of Human Genetics 81:559–575. 10.1086/519795

Robert C, Alves Monteiro HJ, Le Moan A, et al (2025) Fine Scale Patterns of Population Structure and Connectivity in Scandinavian Flat Oysters in Scandinavia (*Ostrea edulis* L.). Evolutionary Applications 18:e70096. 10.1111/eva.70096

Rumberger C, Armstrong M, Kim M, et al (2025) Selection Over Small and Large Spatial Scales in the Face of High Gene Flow. Molecular Ecology 34:e17700. 10.1111/mec.17700

Ryder OA (1986) Species conservation and systematics: the dilemma of subspecies. Trends in Ecology & Evolution 1:9–10. 10.1016/0169-5347(86)90059-5

Saavedra C, Zapata C, Alvarez G (1995) Geographical patterns of variability at allozyme loci in the European oyster *Ostrea edulis*. Marine Biology 122:95–104. 10.1007/BF00349282

Sambade IM, Casanova A, Blanco A, et al (2022) A single genomic region involving a putative chromosome rearrangement in flat oyster (*Ostrea edulis*) is associated with differential host resilience to the parasite *Bonamia ostreae*. Evolutionary Applications 15:1408–1422. 10.1111/eva.13446

Sas H, Deden B, Kamermans P, et al (2020) Bonamia infection in native oysters (*Ostrea edulis*) in relation to European restoration projects. Aquatic Conservation: Marine and Freshwater Ecosystems 30:2150–2162. 10.1002/aqc.3430

Smith RS, Cheng SL, Castorani MCN (2023) Meta-analysis of ecosystem services associated with oyster restoration. Conservation Biology 37:e13966. 10.1111/cobi.13966

Smith RS, Pruett JL (2025) Oyster Restoration to Recover Ecosystem Services. Annual Review of Marine Science 17:83–113. 10.1146/annurev-marine-040423-023007

Tarasov A, Vilella AJ, Cuppen E, et al (2015) Sambamba: fast processing of NGS alignment formats. Bioinformatics 31:2032–2034. 10.1093/bioinformatics/btv098

Thurstan RH, McCormick H, Preston J, et al (2024) Records reveal the vast historical extent of European oyster reef ecosystems. Nature Sustainability 7:1719–1729. 10.1038/s41893-024-01441-4

Triay C, Boizet A, Fragoso C, et al (2025) Fast and accurate imputation of genotypes from noisy low-coverage sequencing data in bi-parental populations. PLOS ONE 20:e0314759. 10.1371/journal.pone.0314759

Valk S, Megens H-J, Kamermans P, Tonk L (2026b) Insights into a native oyster restoration seascape: Genetic divergence between wild, foreign, and cultured *Ostrea edulis* populations. Conservation Genetics 27:56. 10.1007/s10592-026-01758-x

Valk S, Megens H-J, Nijland R, Gundappa MK (2026a) The structural variation landscape of the flat oyster genome. Genomics 118:111274. 10.1016/j.ygeno.2026.111274

Vasimuddin Md, Sanchit Misra, Heng Li, et al (2019). Efficient Architecture-Aware Acceleration of BWA-MEM for Multicore Systems. arXiv:1907.12931v1; 10.48550/arXiv.1907.12931

van Oosterhout C, DeWoody JA, Godoy JA, et al (2025) Genetic load in Evolutionarily Significant Units (ESUs): conservation and management implications. EcoEvoRxiv. 10.32942/X2WD2K

Vera M, Carlsson J, Carlsson JE, et al (2016) Current genetic status, temporal stability and structure of the remnant wild European flat oyster populations: conservation and restoring implications. Marine Biology 163:239. 10.1007/s00227-016-3012-x

Vera M, Pardo BG, Cao A, et al (2019) Signatures of selection for bonamiosis resistance in European flat oyster (*Ostrea edulis*): New genomic tools for breeding programs and management of natural resources. Evolutionary Applications 12:1781–1796. 10.1111/eva.12832

Wang J (2019) A parsimony estimator of the number of populations from a STRUCTURE-like analysis. Molecular Ecology Resources 19:970–981. 10.1111/1755-0998.13000

Whitlock MC, Lotterhos KE (2015) Reliable Detection of Loci Responsible for Local Adaptation: Inference of a Null Model through Trimming the Distribution of F(ST). Am Nat 186 Suppl 1:S24–36. 10.1086/682949

Wickham H (2016). ggplot2: Elegant Graphics for Data Analysis. Springer–Verlag New York. ISBN 978-3-319-24277-4. 260 pp

Yu G (2025). scatterpie: Scatter Pie Plot. R package version 0.2.6, https://github.com/yulab-smu/scatterpie.

Zheng X, Levine D, Shen J, et al (2012) A high-performance computing toolset for relatedness and principal component analysis of SNP data. Bioinformatics 28:3326–3328. 10.1093/bioinformatics/bts606

zu Ermgassen PSE, Spalding MD, Blake B, et al (2012) Historical ecology with real numbers: past and present extent and biomass of an imperilled estuarine habitat. Proceedings of the Royal Society B Biological Sciences 279:3393–3400. 10.1098/rspb.2012.0313

zu Ermgassen PSE, Bos O, Debney, et al. (eds) (2021). European Native Oyster Habitat Restoration Monitoring Handbook. The Zoological Society of London, UK., London, UK. https://archimer.ifremer.fr/doc/00745/85667/

zu Ermgassen PSE, McCormick H, Debney A, et al (2025) European Native Oyster Reef Ecosystems Are Universally Collapsed. Conservation Letters 18:e13068. 10.1111/conl.13068

