## Supplementary Material for "A cryptic local genetic cluster in Northern France amid the European mosaic of flat oyster lineages revealed by integrating SNP array and whole-genome sequencing"

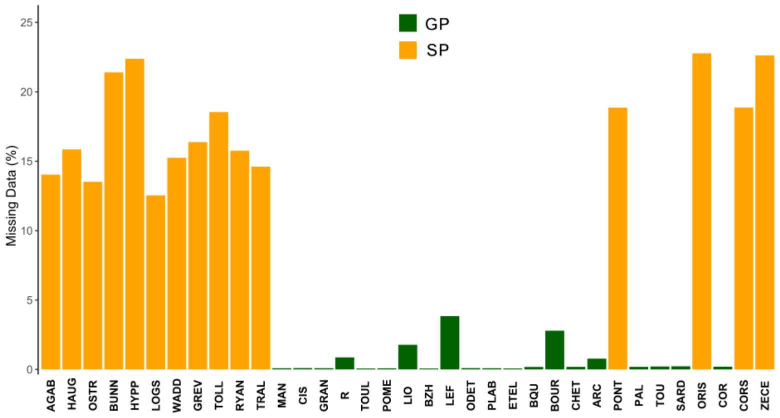


**Fig. S1** Percentage of missing genotypes for the GP and SP of the Merged Dataset II.

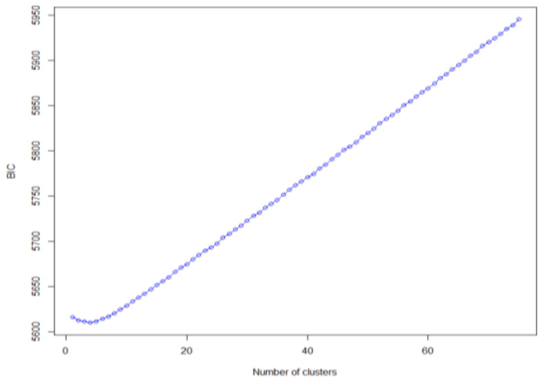


**Fig. S2** Calculation of the best number of clusters through the BIC (Bayesian Information Criterion) of the DAPC analysis.

**Table S1** Values obtained by the three *k*-estimation methods with the best estimate in bold and the next closest one underlined.

| K | Mean_Ln(D/K) | DeltaK | Parsimony |
|---|---|---|---|
| 1 | -548971.52 | - | 0.5000 |
| 2 | -544719.37 | 0.3546 | 0.5217 |
| 3 | -540617.83 | 64.4108 | 0.8403 |
| 4 | -538263.47 | 32.5433 | 0.8841 |
| 5 | -536874.64 | 15.3348 | 0.9000 |
| **6** | **-536124.11** | **69.6680** | **0.9097** |
| 7 | -538096.90 | 3.3338 | 0.8356 |
| 8 | -536688.04 | 8.9842 | 0.9005 |
| 9 | -550235.97 | 0.8144 | 0.8000 |
| 10 | -538143.72 | - | 0.8014 |

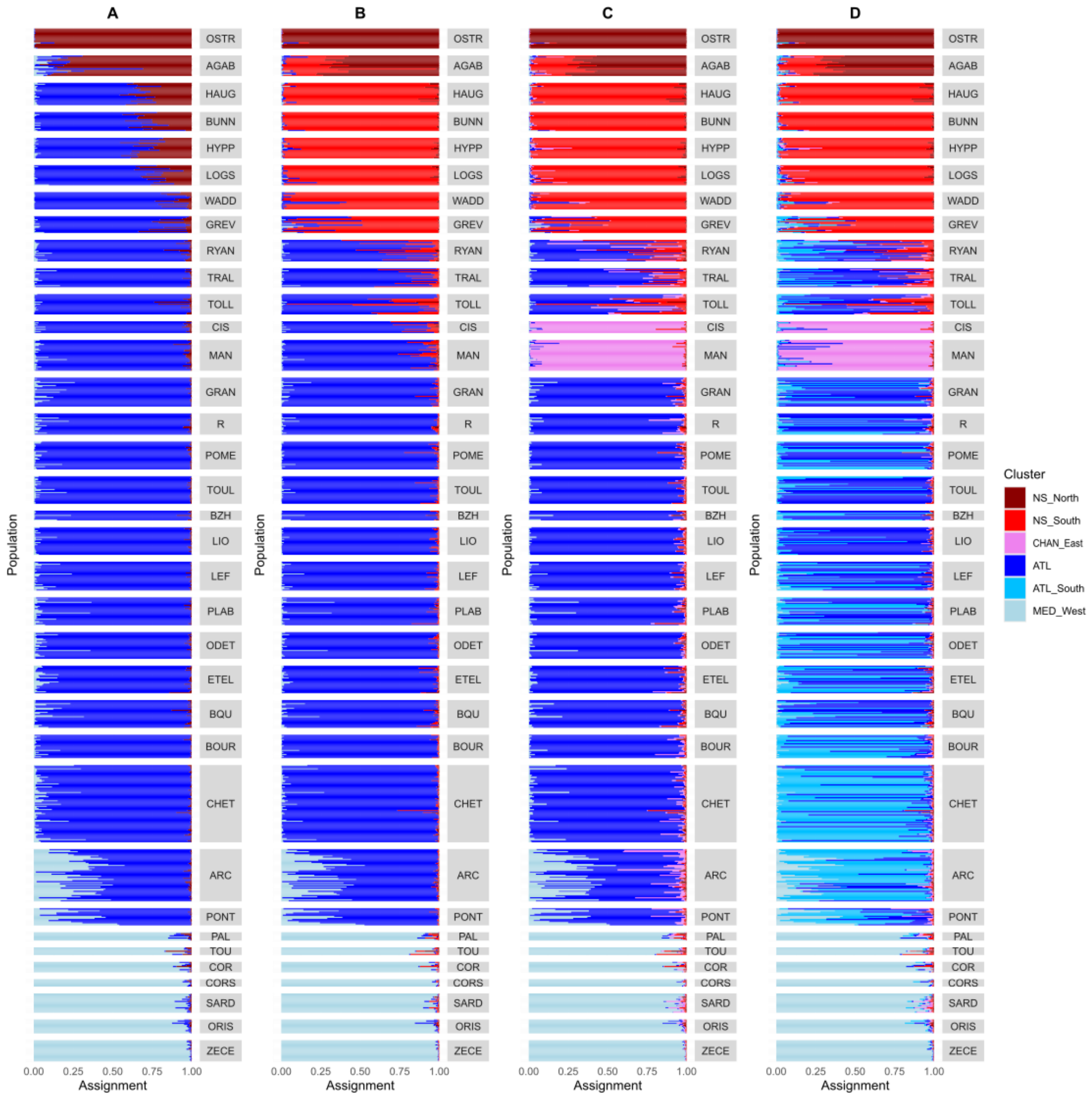


**Fig S3** STRUCTURE analyses indicating individual probability of assignment to different clusters: *k*=3 (A), *k*=4 (B), *k*=5 (C), and *k*=6 (D).

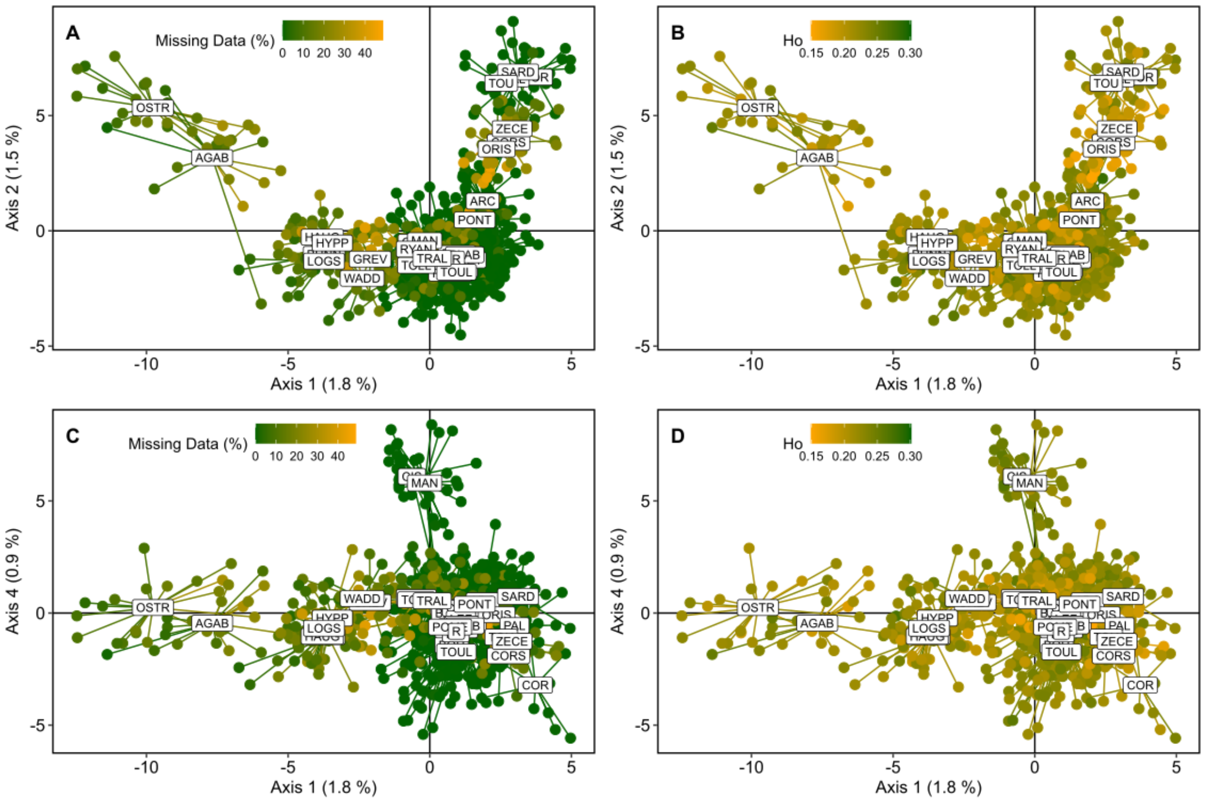


**Fig. S4** PCA representation of axes 1 and 2 (A and B), and axes 1 and 4 (C and D) with the percentage of missing data (A and C) and *Ho* (B and D) indicated by the color gradient.

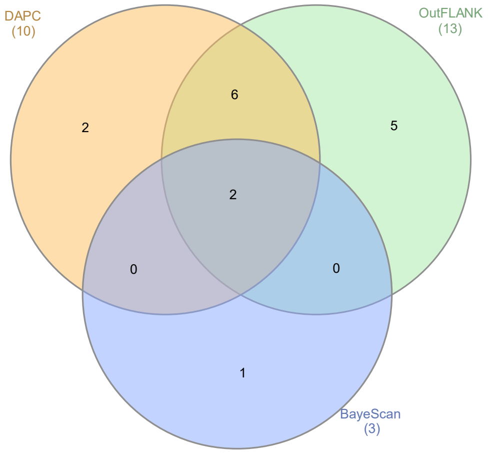


**Fig. S5** Venn diagram of the number of outliers loci obtained with the three methods used.

**Table S2** Outlier loci identified between CHAN_East cluster and all other populations. Index = index of the SNP in the SNP array; Chromosome and chromosome number are from Gundappa et al. (2022); Ni = Number of genes impacted.

| SNP ID | Index | Chromosome | Chromosome number | Sequence position | SNP position | Sequence with [SNP] position | Ni | Genes name |
|---|---|---|---|---|---|---|---|---|
| AX_169159993 | 1251 | NC_079164.1 | 1 | 89592319 | 89592354 | CATGAATCTTTCTGTTCTATGAGTATCAGAGTGTT[A/G]AATGCTGTTTCATGATGCTCTTCACATTCAGAGTC | 1 | trichohyalin-like |
| AX_169172641 | 5634 | NC_079165.1 | 2 | 11984044 | 11984079 | ATATACAAATGTATCATACATTACATAAAACTACA[C/T]GCAGCTTTCATGAAAACAGATTACTCAAGAAAAATGCCCGGATCTAGGTTCCGTCTGGACTCTCTAATAC | 2 | cell division control protein 6 homolog ; NEDD8-conjugating enzyme UBE2F-like |
| AX_169167418 | 3795 | NC_079165.1 | 2 | 12301658 | 12301693 | TAGCAGGTACTAGTCAATGCTGATAGTGGAATCTC[A/G]TTATTGCATCAAAGTTAGCTTTGGGTGAATTCAAT | 1 | catenin beta-like, transcript variant X1 |
| AX_169183303 | 9392 | NC_079165.1 | 2 | 85550598 | 85550633 | CATAACTGAACTCCTACGAAGAAATGCTTTTATTA[A/C]CTTATTCACATTTATGCTCAAAATTTGCCACAGGC | 1 | disco-interacting protein 2 homolog C-like |
| AX_169188101 | 11074 | NC_079165.1 | 2 | 91700881 | 91700916 | TATTCAGAAATTTGCTGTACTTCTGGTAGATAGAA[A/C]TAAAGAGCTCAACTATGAGAAGAAAAAGACGGACT | 1 | uncharacterized LOC125679706 |
| AX_169178760 | 7800 | NC_079166.1 | 3 | 29483198 | 29483233 | TAAGAAAAATCATCTTCGATAGTCAGGTAGGTAAA[G/T]GTTACGATCACCTCTCATCTTGTGAATATTGATTG | 1 | uncharacterized LOC125677342 |
| AX_169183089 | 9320 | NC_079166.1 | 3 | 97147437 | 97147472 | AATGTGAACTAAGTGCGTTTTAGCTAAATAGTCAT[A/G]TTTACTGATTGGTTAATATCATATGCTATCCGAAA | 0 | intergenic |
| AX_169168227 | 4077 | NC_079167.1 | 4 | 33395478 | 33395513 | TGGAGTTAAGGTACTTCCTGGACTTTTACAGTGCG[A/G]ACACTAGAATTTCAATGGAATAAAATATCAAAATA | 0 | intergenic |
| AX_169192390 | 11954 | NC_079167.1 | 4 | 57280206 | 57280241 | TTTTGTAGACATATCATATCGTCTGTCAAAAGCTC[A/G]TGAATTAATTGTTGCGTGCACAAACATACTATTCT | 1 | orexin receptor type 2-like |
| AX_169191970 | 11868 | NC_079168.1 | 5 | 41881671 | 41881706 | CACATTTTAACAATCCATTTCTTTAGATTCCATTT[A/C]TGTTGTAATACTTAATTTCTTTGATCAGATACACA | 2 | delta(14)-sterol reductase LBR-like ; polypeptide N-acetylgalactosaminyltransferase 2-like |
| AX_169191822 | 11842 | NC_079169.1 | 6 | 67772718 | 67772753 | TGCAGATCCATTACTTATAAAAACCTCTGACTAAG[A/G]CACAGAAAATCTTCTCATGGTCACTGTATTCTTAT | 1 | uncharacterized LOC125645948 |
| AX_169158880 | 839 | NC_079170.1 | 7 | 7237099 | 7237134 | TGTTATATGTTATAGGGAAAGCATTTTACACAGCA[A/G]TTGTAAATATCTACGATCAATCATCATTCAAAAAT | 1 | TBC1 domain family member 30-like, transcript variant X2 |
| AX_169171396 | 5168 | NC_079170.1 | 7 | 10434126 | 10434161 | CACCTCAGGTCAGCTGTCAATAATGGTTAACTGTG[C/T]GGCTGTGTGAAAGGGCTGACATATCATTCAGCAAC | 2 | alpha-L-fucosidase-like ; uncharacterized LOC125654012 |
| AX_169172319 | 5502 | NC_079170.1 | 7 | 58038930 | 58038965 | AATGACATTTTGCTGCTCTCTGTCGGACTGGACAA[A/G]CGAATTATTTGCTTTGATGTCCAAAGCAAAAAGTA | 1 | protein NEDD1-like |
| AX_169159375 | 1026 | NC_079171.1 | 8 | 45482133 | 45482168 | ATGACCAGTGATGATGAAAGAAAATCCATAGTTGC[A/G]CGTTACTCTGGTTACACAAAGAAACAAAGCATCCA | 1 | uncharacterized LOC125661341 |
| AX_169156651 | 67 | NC_079172.1 | 9 | 5424589 | 5424624 | AAAGCAAATAGTTCTCATTATTCCTTACAGTTGTT[A/G]GTATTGTATACATTTAATAACGATAAGAACAAATG | 1 | G-protein coupled receptor GRL101-like |
